# Deletion of *Letmd1* leads to the disruption of mitochondrial function in brown adipose tissue

**DOI:** 10.1101/2020.11.18.388199

**Authors:** Runjie Song, Yaqi Du, Peng Li, Huijiao Liu, Han Zheng, Xiaohui Lu, Shenghong Wang, Lijun Zhou, Nafis A Rahman, Sławomir Wołczyński, Adam Kretowski, Fazheng Ren, Xiru Li, Xiangdong Li

**Author notes:** Runjie Song, Yaqi Du, Peng Li, contributed equally to this work. Corresponding author: Fazheng Ren^5^, Xiru Li^6^, Xiangdong Li^1^.

## Abstract

Human cervical cancer oncogene (HCCR-1), also named as LETMD1, is a LETM-domain containing outer mitochondrial membrane protein which plays an important role in the carcinogenesis of cancers. Surprisingly, we found that loss of *Letmd1* in mice leads to multiply severe abnormities, such as the brown adipose tissue (BAT) whitening, disruption of thermogenesis, cold-induced death, diet-induced obesity, hyperglycinemia and insulin resistance. Mechanistically, deletion of *Letmd1* in BAT causes the reduction of mitochondrial calcium ion, which in turn results in the suppressed fission of mitochondria, and ultimately leads to the depletion of *Ucp1*-mediated BAT heat production. This study indicates that LETMD1 plays a crucial role in controlling BAT thermogenesis and energy homeostasis by regulating mitochondrial structures and functions, and also provides a novel insight for the clinical biomarker and therapeutical target of oncogene for the metabolic disorders.

**Highlights:**

1. *Letmd1* is an oncogene and also highly expressed in brown adipose tissue (BAT) of human and mice.
2. Loss of *Letmd1* leads to BAT whitening, diet-induced obesity, hyperglycemia and insulin resistant.
3. *Letmd1* knockout causes the disruption of thermogenesis and death at 4°C exposure.
4. Deletion of *Letmd1* results in mitochondrial calcium homeostasis disorders.

**Graphic abstract:** 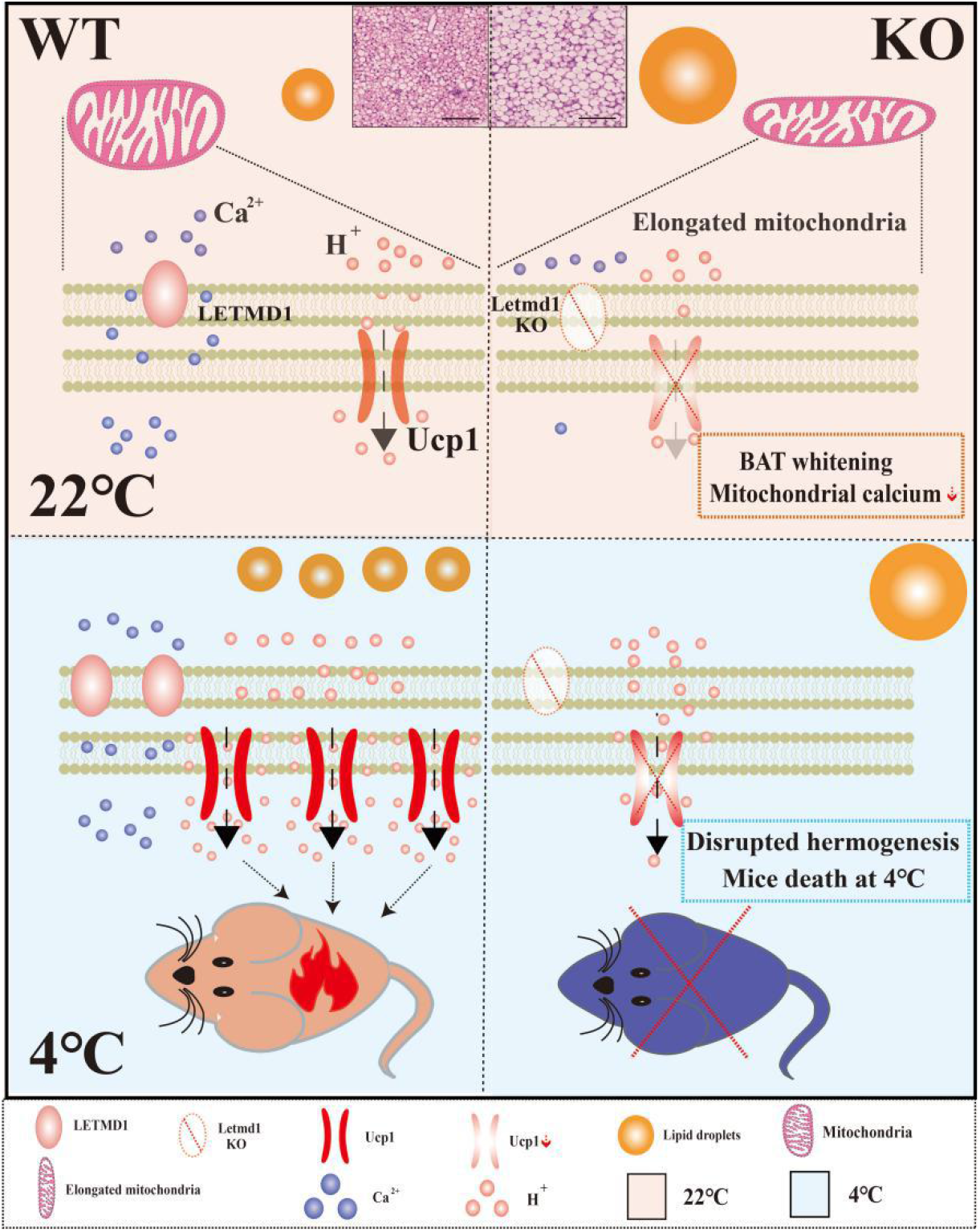

## 1. Introduction

Leucine zipper-, EF-hand-containing transmembrane protein 1 domain containing 1 (LETMD1), also called human cervical cancer oncogene (HCCR-1) is located in the outer membrane of mitochondria^[1]^. Previous studies have shown that *LETMD1* served as an oncogene and highly expressed in various malignant tumors by regulating phosphatidylinositol-4,5-bisphosphate 3-kinase (PI3K)/serine/threonine kinase (Akt) or P53 signal pathway, such as cervical cancer, breast cancer, hepatocellular carcinoma (HCC), gastric cancer and colon cancer^[2–8]^. Furthermore, the high expression of LETMD1 could be served as an early clinical diagnosis bio-marker for HCC and breast cancer^[6, 9]^. A recent study firstly mentioned the regulation of LETMD1 in mitochondrial hyperpolarization^[10]^.

Leucine zipper-, EF-hand-containing transmembrane protein 1 (LETM1), a homologous gene of LETMD1 is an inner membrane protein of mitochondria^[11]^. LETM1 was identified as a key component of ions transporter in mitochondria, including Ca^2+^/H^+^ transportation, K^+^/H^+^ exchange, Na^+^/Ca^2+^ exchange, and Mg^2+^ transportation to maintain the mitochondrial biology and cation homeostasis^[12–16]^. Most studies have shown that knockdown of *LETM1* caused the mitochondrial swelling in worms, drosophila, yeast, and protists^[17–19]^. In addition, previous studies have shown that the mutation of four amino acid residues in the LETM-domain resulted in the abnormal structure in mitochondria of yeast and LETM-domain was highly conserved in yeast, worms, drosophila, plants and mammals^[20–22]^. The fundamental functions of *Letm1* resulted in the gene knockout animals of it was unavailable^[23]^, which hampers the better understanding of the real physiological function of *LETM1*.

Interestingly, by analysis the expression profiles of *LETMD1* in human and mice, we found that *LETMD1* was highly expressed in the metabolism relative tissues, especially brown adipose tissue (BAT), and the expression of *LETMD1* was significantly reduced in adipose tissues of the obese people and the high fat diet (HFD) induced mice.

Some genes located on the mitochondrial membrane are critical to the integrity of the mitochondrial structure, including uncoupling protein 1 (*Ucp1*), mitofusin 2 (*Mfn2*), metalloprotease 1 (*Oma1*), apoptosis inducing factor mitochondria associated 2 (*Aifm2*)^[24–27]^. The deletion of these genes can cause abnormal mitochondrial morphology, which leads to the abnormal mitochondrial function. These genes have been reported to be related to the functions of BAT or beige fat^[24–27]^. BAT loads more mitochondria in order to process mitochondrial respiration to maintain its normal physiological function of thermogenesis^[28]^. It has been reported that dysfunctions of BAT were related with the pathophysiology of a variety of metabolic diseases, such as type 2 diabetes, obesity, dyslipidemia, and cardiovascular diseases^[29–32]^.

Therefore, we hypothesis that LETMD1 may play an important role in the metabolism of BAT through regulating mitochondrial functions. To test our hypothesis, we generated the *Letmd1* knockout (KO) mice and used them to address the critical roles of *Ledmd1* in the whitening of BAT, thermogenesis, and glucose intolerance and insulin resistance. Mechanically, our findings outline an important intrinsic role for LETMD1 in the fissions of mitochondria and Ca^2+^ homoeostasis in mice.

## 2. Materials and Methods

### 2.1 Chemicals

DMEM, glucose, HEPES, PBS, RIPA buffer, enzyme chemiluminescence reagent, insulin, norepinephrine (NE), triiodothyronine (T3), 3-isobutyl-1-methylxanthine (IBMX), dexamethasone (DEX), penicillin-streptomycin (Pen/Strep), CL316,243 (the β-adrenergic receptor agonist), isoflurane, Triton X-100, type I collagenase, NaCl, Na_3_PO_4_, KCl, MgSO_4_ and trypsin were purchased from Sigma, USA. SYBR Green Master Mix Reagent was purchased from Roche, Switzerland. Hoechst 33342 was purchased from Beyotime, China. TRIzol reagent was purchased from Invitrogen, USA. Fetal bovine serum (FBS) was purchased from Gibco, USA. XF assay medium, 2-deoxy-D-glucose (2-DG), Oligomycin, carbonyl cyanide 4-(trifluoromethoxy) phenylhydrazone (FCCP), antimycin A and rotenone were purchased from Agilent, USA. MitoTracker-mitochondrion-selective probes, moloney murine leukemia virus (M-MLV) reverse transcriptase, oligo (dT) and rhod-2 were purchased from Thermo scientific, USA.

### 2.2 Animals

C57BL/6J mice were obtained from Beijing Vital River Laboratory Animal Technology Co., Ltd. China, and were housed with controlled light (12 h light and 12 h in darkness) and temperature (22°C ± 1). Food and water were taken *ad lib*. For the cold challenge, mice were transferred to 4°C, the controls were kept at 22°C, and the body temperature of each mouse was monitored every 2 h for a total of 6-8 h of period. For β-adrenergic receptor agonist treatment, CL316,243 was dissolved in PBS and injected to the mouse intraperitoneally once daily. The animals were injected at a dose of 1 mg/kg for consecutive 10 d. All animals were sacrificed by inhalation of carbon dioxide. The tissues were dissected and quick-frozen for the future analysis, or immediately fixed in 4% paraformaldehyde buffer for the histological study. All the animal experiments were approved by the ethic committee of the China Agricultural University.

*Letmd1* KO mice (C57BL/6 background) were tailor made by Nanjing biomedical research institute of Nanjing University via CRISPR/Cas9 technology. Cas9 mRNA and sgRNA were co-injected into zygotes. SgRNA directed Cas9 endonuclease cleavage in upstream of exon3 and downstream of exon7, and created a double-strand break (DSB). Such breaks were repaired by non-homologous end joining (NHEJ), and resulted in disruption of *Letmd1. Letmd1* KO mice were generated a 3995 bp chromosomal deletion at *Letmd1* locus in the mouse genome. Mice were fed with chow diet (LOT number H10010, HFK bioscience, Beijing) or HFD (LOT number H10060, HFK bioscience, Beijing).

### 2.3 Real-time quantitative PCR (RT-qPCR) analysis

For RT-qPCR analysis, TRIzol reagent was used to extract total RNA from cells and tissues. RNA samples were treated with RNase-free DNase I to remove genomic DNA. The purity and concentration of RNA samples were measured by Nanodrop 2000 (ThermoFisher, USA). Ratios of absorption (260/280 nm) of RNA samples were between 1.8 and 2.0. Then 1 μg of RNA were reversed transcribed using M-MLV reverse transcriptase and oligo (dT). The cDNA was amplified in the Roche Light Cycler 480 system (Roche, Switzerland) with SYBR Green Master Mix Reagent. All data was analyzed by Using LightCycler®480 software. The melting curve was analyzed to confirm the specificity of the signal. The relative gene expression levels in each sample were normalized to the level of β-actin. The RT-qPCR primers used were listed in the Supplementary Table 1.

### 2.4 Western Blot

The Total protein was extracted with RIPA buffer. Protein concentrations were measured using BCA Protein Assay Reagent (Abcam, USA). Proteins were separated on SDS-PAGE and transferred to the polyvinylidene fluoride (PVDF) membrane (Millipore, USA) and blocked in 5% fat-free milk for 45 min at 25°C. After washing, the membrane was incubated with the antibodies and visualized by using an enzyme chemiluminescence reagent. The antibodies used are listed in Supplementary Table 2.

### 2.5 Cell culture

Primary brown adipocytes were isolated from the interscapular BAT depot of the wild-type (WT) and *Letmd1* KO newborn mice. BAT depot was minced and incubated in the DMEM containing 1.5 mg/mL type I collagenase at 37°C 40 min for digestion. The stromal vascular fraction (SVF) was filtered through a mesh filter of 74 μm and plated in DMEM containing 20% FBS and 1% Pen/Strep. A part of SVF was dissociated with trypsin and resuspended with DMEM containing 20% FBS before staining with the following antibodies for 10 min on ice: Sca-1-APC, CD11b-FITC, and CD45-PE. Following antibody incubation, cells were washed, centrifuged at 800 rpm for 10 min, and sorted with a BD FACSAria (BD Biosciences, USA). Data analysis was performed by using BD FACS Diva software and showed primary brown adipocytes (SCA1+/CD31 –/CD11b –) were obtained at 95% purity. Once the cells reach confluence, DMEM (10% FBS, 20 nM insulin, 1 nM T3, 0.5 mM IBMX and 2 μg/mL DEXA) was used to induce the differentiation. Two days later, the medium was replaced every 2 d with DMEM (10%FBS, 20 nM insulin and 1 nM T3) for 6 d.

### 2.6 Blood chemistry measurement

Blood samples were collected by cardiac puncture after anesthesia by isoflurane inhalation. Serum were obtained by centrifuging blood samples at 3000 rpm for 5 min. Serum triglyceride determination (TG) kit (Sigma, USA) and free fatty acid assay (FFA) kit (Abcam, USA) were used to measure the triglyceride and free fatty acid, separately. Ultra-sensitive mouse ELISA kit (Crystal Chem, USA) was used to determine the level of serum insulin.

### 2.7 Glucose and insulin resistance tests

Five μL blood was collected from the tail vein of mice, and dropped into the glucose test strip (Roche, Switzerland) and measured by the glucometer (Roche, Switzerland). For glucose tolerance test (GTT), after the overnight fasting, glucose (2 g/kg of body weight for mice fed chow diet, 1 g/kg for mice fed HFD) was administered intraperitoneally. The blood glucose was measured at 0, 15, 30, 60, 90 and 120 min. For insulin tolerance test (ITT), mice fasted for 4 h were injected with insulin (0.75 U/kg of body weight for mice fed chow diet, 1.5 U/kg for mice fed HFD) intraperitoneally. The blood glucose was measured at 0, 15, 30, 60, 90 and 120 min.

### 2.8 Seahorse assay

Primary brown adipocytes were trypsinized, and reseeded in XF24 plates at 50 K cells per well assayed on day 6 of differentiation (mature brown adipocytes). According to the manufacturer’s instructions, a Seahorse XF24 analyzer (Agilent Technologies, USA) was used to measure the oxygen consumption rate (OCR) and extracellular acidification rate (ECAR) of the cells. Cells were incubated in XF assay medium, supplemented with 25 mM glucose, 1 mM pyruvate for 1 h before the measurement. Basal Respiration were recorded, max respiration was determined by final concentration of 1.5 μM Oligomycin and 1 μM FCCP. The final concentration of 0.5 μM of antimycin A and rotenone and 0.5 μM rotenone were used to inhibit Complex III and Complex I dependent respiration. For the glycolysis stress test, 10 mM glucose,1 μM Oligomycin, and 50 mM 2-DG were sequentially injected. Results were analyzed by the Seahorse XF24 analyzer software.

### 2.9 Mitochondrial content measurement

The total DNA from BAT was extracted by using Quick Extract DNA Extraction Solution 1.0 (Epicenter, USA). RT-qPCR was used to determine the mitochondrial DNA copy number by calculating the ratio of amplification between mitochondrial DNA and nuclear DNA, primers for mitochondrial DNA and nuclear DNA amplification were listed in Supplementary Table 1.

### 2.10 Histology and immunostaining

For histology, the tissues were fixed with 4% paraformaldehyde, embedded in paraffin, and sectioned at 4 μm. After dewaxing, sections were stained with hematoxylin and eosin (H&E). For immunostaining, sections were pretreated with hydrogen peroxide (3%) for 10 min to remove the endogenous peroxidase, followed by antigen retrieval in a microwave for 15 min in 10 mM citrate buffer (pH 6.0). UCP1 primary antibody was used at a dilution of 1:500 and incubated for 30 min at room temperature, followed by washing and incubation with the biotinylated secondary antibody for 30 min at room temperature and coloration with 3,3’-diaminobenzidine (DAB). The slides were counterstained with hematoxylin and dehydrated in alcohol and xylene before mounting.

### 2.11 Mitochondrial morphology detection

Primary brown pre-adipocytes were detected with MitoTracker-mitochondrion-selective probes. Live cell fluorescence images were captured with laser scanning confocal microscope A1 (Nikon, Japan). The nucleus of live cells was stained with Hoechst 33342. The 405 nm laser was used to detect Hoechst. The 488 nm laser was used to detect the activated MitoTracker-mitochondrion-selective probes. Mitochondria from each group were randomly selected.

### 2.12 The concentration of Ca^2+^ measurement

In order to image mitochondrial Ca^2+^, the differentiated brown adipocytes were incubated with 5 μM rhod-2 for 15 min at 37°C, and then were changed to the photo buffer [125 mM NaCl, 5 mM KCl, 1 mM Na_3_PO_4_, 1 mM MgSO_4_, 5.5 mM glucose and 20 mM HEPES (pH 7.4)]. After NE stimulation, a series of images were captured every 10 s using a confocal microscope (Nikon, Japan). The 561 nm laser was used to excite rhod-2. ImageJ (National Institutes of Health, USA) was used to analyze and quantify the recorded images.

### 2.13 Energy balance research

Energy balance parameters of the animals were determined in a computer-controlled open circuit system (Oxymax, USA), which is part of the integration comprehensive laboratory animal monitoring system (Columbus Instruments, USA), the O_2_ consumption, CO_2_ production, food intake, Respiratory Quotient (RQ), Energy exchange rate (EE) and the activity level were monitored by the CLAMS system (Columbus Instruments, USA).

### 2.14 Metabolic parameter measurement

Fat and lean mass were measured by the nuclear magnetic resonance spectrometer (Shanghai electronic technology co., Ltd, China). The infrared video was measured by an infrared thermal imager (Magnity Electronics Co.,Ltd, China).

### 2.15 Bioinformatics analysis

The gene microarray data (GSE8044, GSE1342) was downloaded from Gene Expression Omnibus (GEO). The GSE129084, GSE72603, GSE113764 and 32 cancer datasets were download from GEO and TCGA (The Cancer Genome Atlas) database in FPKM (Fragments Per Kilobase Million) or RPKM (Reads Per Kilobase Million) format and transferred to TPM (Transcripts per million reads). The edgeR or DEseq2 was used to screen the differentially expressed genes (DEGs). Gene Set Enrichment Analysis (GSEA) analysis was performed to show the unique biological significance. The GSEA analysis were carried out according to the protocol (http://www.gsea-msigdb.org/gsea/)^[33]^. The ggplot2 package in R was used to show the pathway maps.

### 2.16 Statistical analysis

All analyses were performed at least in triplicate, and the means obtained were subjected to Unpaired two-tailed Student’s t-tests. In the figures, asterisks denote statistical significance (*P < 0.05, **P < 0.01, ***P < 0.001). All data were reported as the mean ± standard error of mean (SEM). The means and SEM from at least three independent experiments were presented in graphs.

## 3 RESULTS

### 3.1 Letmd1 is highly expressed in BAT both in human and mice

Previous studies have showed that LETMD1 was conserved in mitochondrial outer membrane^[1]^, and served as an oncogene to participate in various of cancers^[2]^. The mitochondria location of LETMD1 indicates that it might play a role in metabolism. By mining the data and re-analyzed the GSEA enrichment from both the TCGA database and hBAT (human brown adipose tissue) database (GSE113764), we found that *LETMD1* was positively correlated with ATP metabolic process and fatty acid metabolism (Fig. 1a and b).

**Fig. 1.**
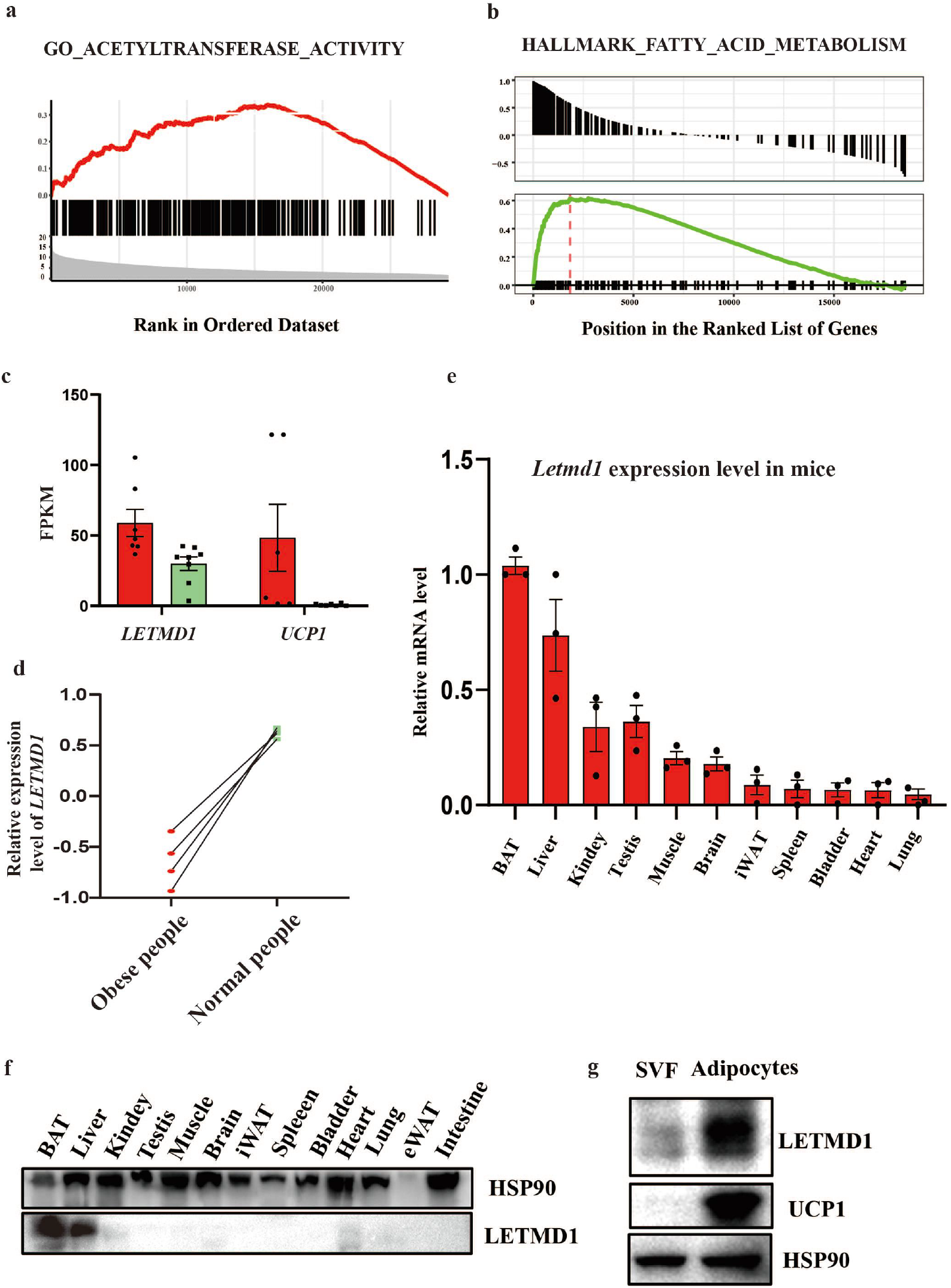
*LETMD1* is highly expressed in BAT of mice and human. (a-b). *LETMD1* was correlated positively with gene signatures related to ATP metabolic process and fatty acid metabolism. (c) *LETMD1* FPKM of human BAT and WAT. (d) The relative expression level of *LETMD1* in obese people and normal people. (e) Relative mRNA expression of *Letmd1* in different mice tissues (n = 3) by RT-qPCR, β-actin was used as the internal control. (f) Immunoblotting measurement of LETMD1 in different mice tissues (n = 3), HSP90 was used as a loading control. (g) Representative immunoblotting measurement of LETMD1 in human SVF cells and human adipocytes, HSP90 was used as a loading control. Data were mean ±SEM and each dot showed as one replicate.

Comparing the mRNA levels of hBAT and human white adipose tissue (hWAT) in 9 cases from this hBAT database (GSE113764), the expression level of *LETMD1* in hBAT was higher than that in WAT (Fig. 1c). Moreover, by re-analysis of the transcriptional profiling of subcutaneous adipose tissue of morbidly obese subjects (GSE66921), we found that the expression of *LETMD1* was markedly upregulated when the obese people turned to the normal body mass index by bariatric surgery (Fig. 1d). In addition, we found that the highest expression level of *Letmd1* was in BAT from the WT mice among other tissues (Fig. 1e and f). Similarly, the highly expressed LETMD1 was observed in the differentiated hBAT-SVF cells (Fig. 1g).

### 3.2 The reduced energy consumption and glucose tolerance in Letmd1 KO mice

To further explore the role of *Letmd1* in metabolism, we generated *Letmd1* KO mice. *Letmd1* KO mice were born at the expected Mendelian frequency. The KO efficiency of LETMD1 was determined by Western Blot (Fig. 2a). No significant difference in body weight between *Letmd1* KO and WT mice with chow diet was observed (Fig. 2b), whereas nuclear magnetic resonance spectrometer scanning showed the body fat mass index was increased in KO mice (Fig. 2c). Accordingly, the whole-body OCR of *Letmd1* KO compared to WT mice measured by CLAMS was significantly lower when mice were kept at 22°C (Fig. 2d and e), while no significant differences in locomotor activity levels and the food consumption from the *Letmd1* KO mice were observed (Fig. S1a-d).

**Fig. 2.**
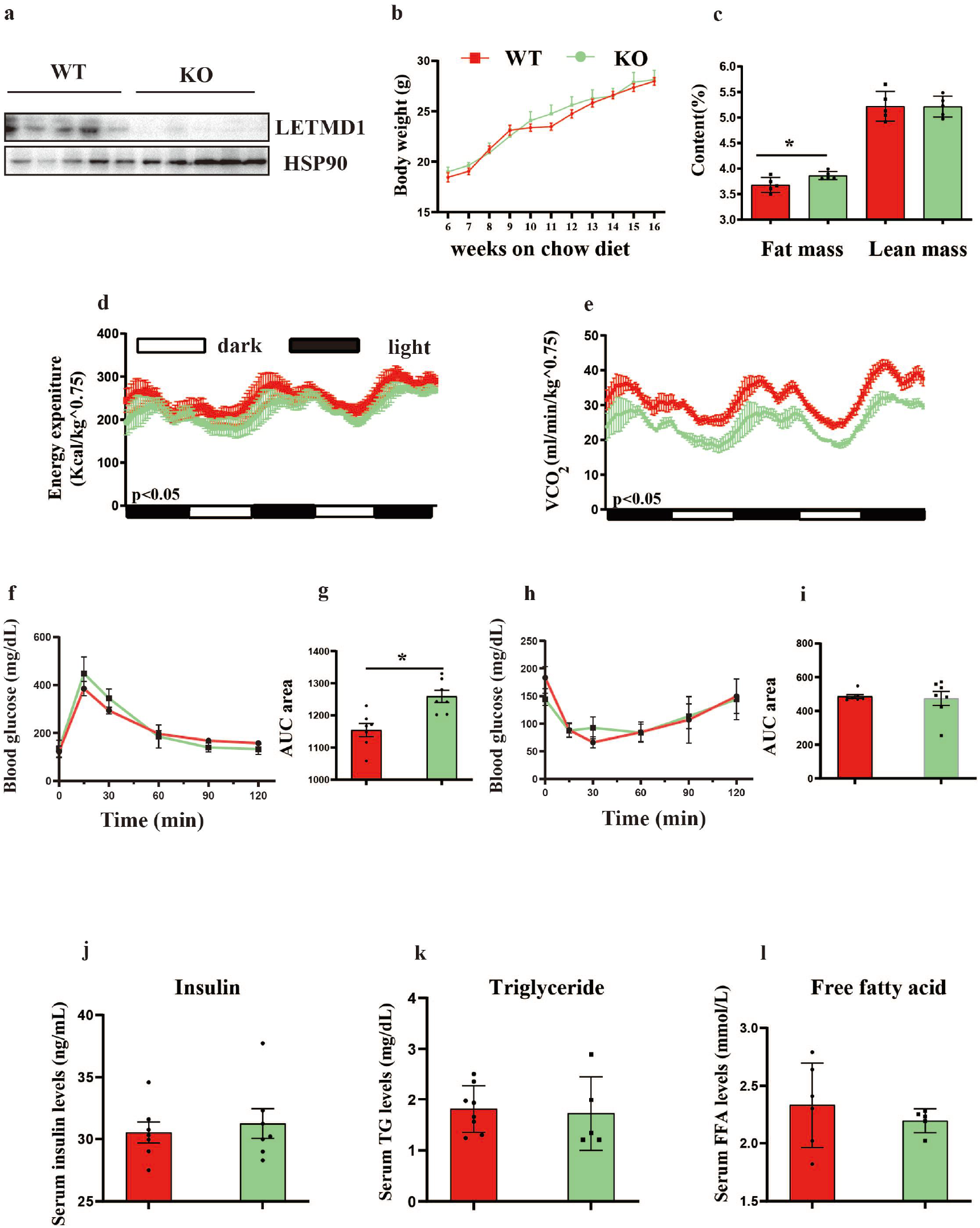
The reduced energy consumption and glucose tolerance in *Letmd1* KO mice. (a) Representative immunoblotting measurement of LETMD1 in BAT of WT and KO mice (n = 5 for each group). (b) Body weight measurements of WT and KO mice fed with chow diet at 22°C (n = 7 for each group). (c) The fat and lean mass of WT and KO mice fed with chow diet at 22°C (n = 5 for each group) measured by the magnetic resonance spectrometer. (d-e) The energy expenditure and O_2_ consumption rates in 8-w-old WT and KO mice at 22°C (n = 5 for each group) measured by the CLAMS system. (f-g) The blood glucose levels with glucose injection (2 g/kg) measured by the glucometer as GTT for WT and KO mice fed with chow diet at 22°C (n = 6 for each group). (h-i) The blood glucose levels with insulin injection (0.75 U/kg) measured by the glucometer as ITT for WT and KO mice fed with chow diet at 22°C (n = 6 for each group). (j-l) Serum levels of Triacylglycerol (TG), Free Fatty Acids (FFA) and Insulin in WT and KO mice fed with chow diet at 22°C (n = 5-8). Data were mean ±SEM and each dot showed as one replicate. *P < 0.05, **P < 0.01, ***P < 0.001, Unpaired two-tailed Student’s t-tests was used in two sets of data.

Furthermore, GTT and ITT showed that the glucose-tolerant of *Letmd1* KO mice was decreased than WT mice with chow diet at 22°C (Fig. 2f-i), but there was no significant difference in insulin sensitivity (Fig. 2h and i). No significant difference in the contents of insulin, free fatty acid and triglyceride between *Letmd1* KO and WT mice were observed (Fig. 2j-l). Taken together, our study demonstrated that deletion of *Letmd1* resulted in reducing of the basal metabolism and the glucose intolerance in mice.

### 3.3 Letmd1 deficiency results in BAT whitening and disruption of mitochondrial function in BAT

The histopathological analysis revealed that BAT of *Letmd1* KO underwent whitening (Fig. 3a), while no significant differences were seen in other fat tissues and liver in *Letmd1* KO mice (Fig. S1e-g). BAT is an energy metabolic pool, which is characterized by abundant mitochondria and unique expression of UCP1 for thermogenesis^[34–36]^. Next, we checked a series of gene expressions which related to thermogenesis, electron transport, fatty acid oxidation and synthesis, glucose oxidation in the BAT of *Letmd1* KO mice. The results showed that the expressions of thermogenesis specific genes, electron transport related genes, fatty acid oxidation and glucose oxidation related genes were decreased compared to the WT mice (Fig. 3b and c).

**Fig. 3.**
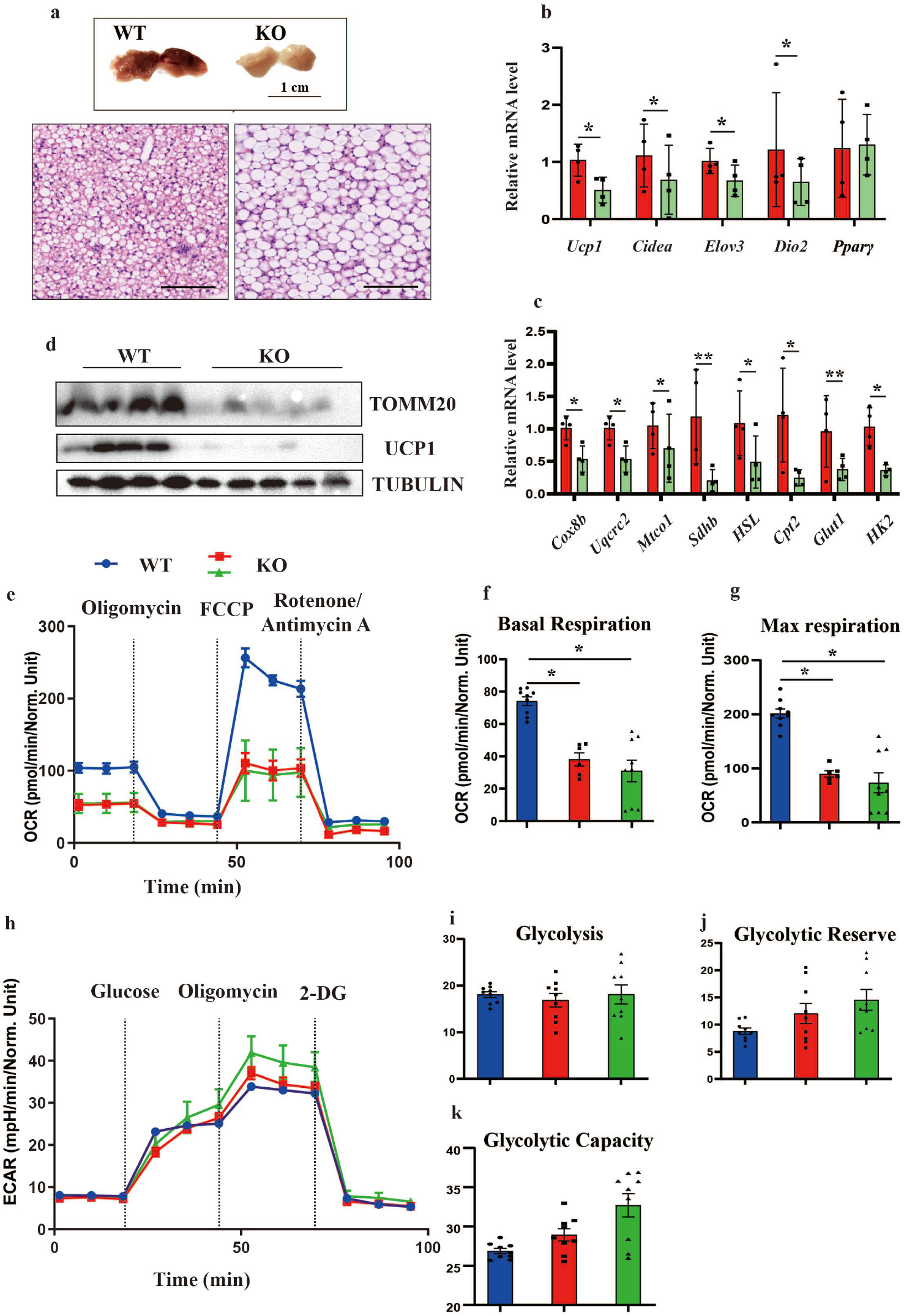
*Letmd1* deficiency results in BAT whitening and disruption of mitochondrial function in BAT. (a) The photograph of BAT in WT and KO mice (Scale bar 1 cm) and representative optical microscopy images of H&E staining BAT in WT and KO mice under chow diet at 22°C (Scale bar 200 μm). (b-c) Relative mRNA expression of thermogenesis, respiratory chain, fatty acid and glucose oxidation related genes in BAT analyzed by RT-qPCR, β-actin was used as the internal control. (n = 4-5 in each group). (d) Immunoblotting measurements of TOMM20 and UCP1 in BAT of WT and KO mice (n = 4-6 in each group), α-TUBULIN was used as a loading control. (e-k) OCR and ECAR measurements by Seahorse XF24 of differentiated brown adipocytes isolated from BAT of WT and KO mice, (f and g) showed basal and maximal respiration rates, (i-k) showed basal glycolysis, glycolytic reserve and glycolytic capacity respectively. Oligomycin, FCCP, and rotenone/antimycin A were added at the time points in OCR test and Glucose, Oligomycin and 2-DG were added at the time points in ECAR test indicated by dashed lines. Data were mean ±SEM and each dot showed as one replicate. *P < 0.05, **P < 0.01, ***P < 0.001, Unpaired two-tailed Student’s t-tests was used in two sets of data.

Furthermore, Western Blot analysis showed that the expressions of UCP1 and TOMM20 were drastically decreased, which indicated that the mitochondrial functions might be destroyed in BAT of *Letmd1* KO mice (Fig. 3d). In order to determine the effects of *Letmd1* on the mitochondrial capacities in BAT, cell metabolism analysis was performed on the differentiated pre-adipocytes isolated from the *Letmd1* KO and WT mice.

Compared with the WT group, the OCR (the capabilities of basal respiration and the max respiration) were significantly reduced in *Letmd1* KO group (Fig. 3e-g). Next, the maximal glycolysis flux capacities of the *Letmd1* KO group measured by extracellular acidification rate (ECAR) was no significant differences (Fig. 3h-k).

### 3.4 Letmd1 is critical for thermogenesis in BAT

As BAT is the main tissue for thermogenesis, and deletion of *Letmd1* caused the dysfunctions of mitochondria which might result in a defeat for maintaining the homoeostasis of body temperature by BAT thermogenesis to resist the cold environment.

To unveil the relationship between *Letmd1* and BAT thermogenesis, we mined and re-analyzed the database (GSE72603, GSE13432, GSE8044 and GSE129083), we found that *Letmd1* and other thermogenic genes were up-regulated after cold exposure and CL316,243 injection (Fig. 4a). Consistent with these analyzed results, we found that the expression of *Letmd1* was up-regulated which has the same expression tendency with *Ucp1* after 7 d cold exposure (4°C) or CL316,243 injection (Fig. 4b and c), respectively. These results indicated that *Letmd1* might participate in the thermogenesis. Subsequently, *Letmd1* KO and WT mice were exposed to the acute cold environment (4°C) to test the thermogenesis. Surprisingly, *Letmd1* KO mice showed cold intolerance with the dramatic decreases of the core body temperatures compared to WT mice, and KO mice were died between 6-8 h within the cold exposure, but WT mice could maintain the body temperature to resist the cold condition (Fig. 4d and e).

**Fig. 4.**
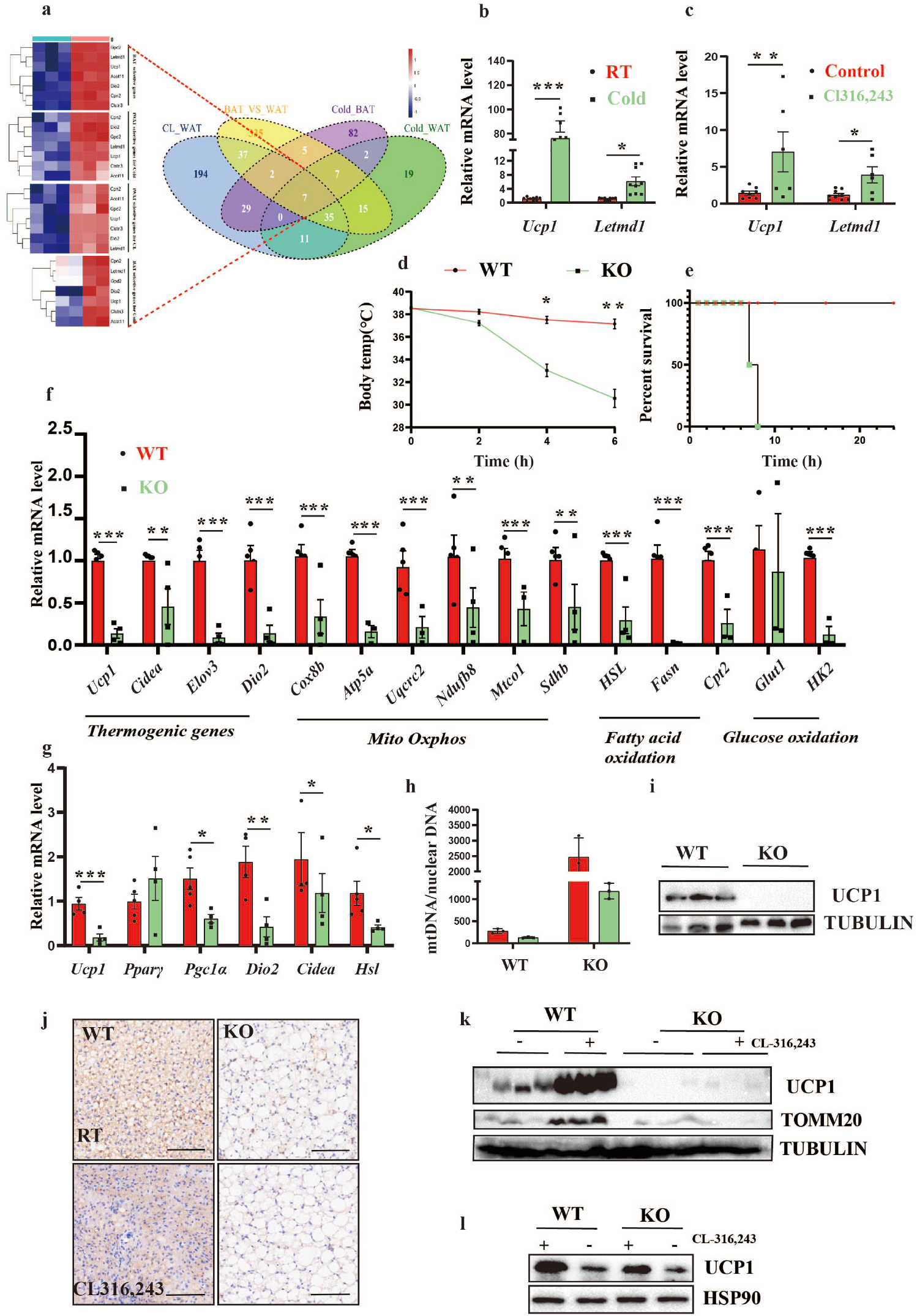
*Letmd1* is critical for thermogenesis in BAT. (a) Veen diagram of search strategy to identify factors highly expressed by *Ucp1* in adipose tissue. The following publicly available datasets were used for the comparisons: BAT and WAT following 1 or 5 w cold exposure (GSE72603 and GSE13432), BAT versus iWAT (GSE8044). WAT following 5 d following cold exposure (GSE129083). (b-c) Relative mRNA expression of *Ucp1* and *Letmd1* in BAT following 7 d cold exposure or 10 d CL316,243 treatment analyzed by RT-qPCR, β-actin was used as the internal control. (n =4-7 for each group). (d) The core temperature was tested by the thermo detector every 2 h of WT and KO mice in acute cold exposure (4°C) (n = 6-8 for each group). (e) Kaplan-Meier survival curves of WT and KO mice in acute cold exposure (4°C) (n = 7-8 for each group). (f) Relative mRNA expression of thermogenesis, respiratory chain, fatty acid and glucose oxidation related genes in BAT of WT and KO mice following 6 h cold exposure analyzed by RT-qPCR, β-actin was used as the internal control. (n = 4-5 in each group). (g) Relative mRNA expression of thermogenesis related genes in BAT of WT and KO mice following 10 d CL316,243 analyzed by RT-qPCR, β-actin was used as the internal control. (n = 4-5 in each group). (h) mtDNA copy number normalized to nuclear DNA was measured by RT-qPCR in BAT of WT and KO mice at 22°C or 4°C exposure (n =3-4 in each group). (i) Immunoblotting measurements of TOMM20 and UCP1 in BAT of WT and KO mice following 6 h cold exposure (n=3 in each group), α-TUBULIN was used as a loading control. (j) Representative optical microscopy images of UCP1-IHC staining BAT in WT and KO mice with or without CL316,243 treatment (Scale bar 200 μm). (k) Immunoblotting measurements of UCP1 and TOMM20 from BAT of WT and KO mice with or without CL316,243 treatment (n = 3-4 in each group), α-TUBULIN as loading control. (l) Representative immunoblotting measurement of UCP1 in differentiated brown adipocyte isolated from WT and KO mice with or without 6 h CL316,243 treatment. HSP90 was used as a loading control. Data were mean ±SEM and each dot showed as one replicate. *P < 0.05, **P < 0.01, ***P < 0.001, Unpaired two-tailed Student’s t-tests was used in two sets of data.

Furthermore, we found that the expression of the thermogenesis related genes, electron transport and fatty acid oxidation synthesis related genes were significantly down regulated in BAT of Letmd1 KO mice under the cold and CL316,243 treatments compared to WT mice (Fig. 4f and g). Moreover, we detected that the copy number of mitochondrial DNA (mtDNA) was significantly reduced in BAT of Letmd1 KO mice both at the room temperature and the cold treatment (Fig. 4h). In addition, the expression level of Ucp1 was also dramatically decreased in the BAT of Letmd1 KO mice (Fig. 4i).

Western Blot analysis and IHC confirmed that the expression of UCP1 was significantly down regulated in BAT of Letmd1 KO mice at the room temperature and CL316,243 treatment compared to WT mice, respectively (Fig. 4j and k). Moreover, the low expression of TOMM20 in BAT of *Letmd1* KO mice was observed (Fig. 4k).

In order to confirm that the decrease expression of UCP1 was a cellular autonomic response, we treated the differentiated brown adipocytes isolated from *Letmd1* KO mice with CL316,243 for 6 h, and found that the expression of UCP1 was lower than that in WT mice (Fig. 4l). These data indicated that LETMD1 was essential for UCP1 mediated mitochondrial uncoupling thermogenesis in BAT.

### 3.5 Letmd1 deficiency leads to diet induced obesity and insulin resistance

The dysfunctions of BAT can induce obesity and insulin resistance^[37, 38]^. To test the role of *Letmd1* in obesity, WT and *Letmd1* KO mice were fed with HFD for 30 w at 22°C. After 1 2 w HFD feeding, the body weight of *Letmd1* KO mice was higher than WT mice (Fig. 5a and b). Anatomic fat pool showed that weights of BAT, inguinal white adipose tissue (iWAT) and epididymal white adipose tissue (eWAT) were increased in *Letmd1* KO mice fed with HFD (Fig. 5c), while no significant differences in liver weight were found in *Letmd1* KO and WT group (Fig. 5c). The whole-body OCR of *Letmd1* KO compared to WT mice measured by CLAMS was significantly lower when mice were kept at HFD (Fig. 5d and e, Fig. S1h and i), while no significant differences in the food consumption from the *Letmd1* mice were obtained (Fig. S1j). Histological analysis showed that BAT whitening in *Letmd1* KO mice was more severe after 30 w of HFD (Fig. 5f). The results of GTT and ITT showed that *Letmd1* KO mice had the compromised glucose tolerance and insulin sensitivity than WT mice (Fig. 5g and h). No significant changes in free fatty acid and triglyceride in *Letmd1* KO mice were observed (Fig. 5i-k).

**Fig. 5.**
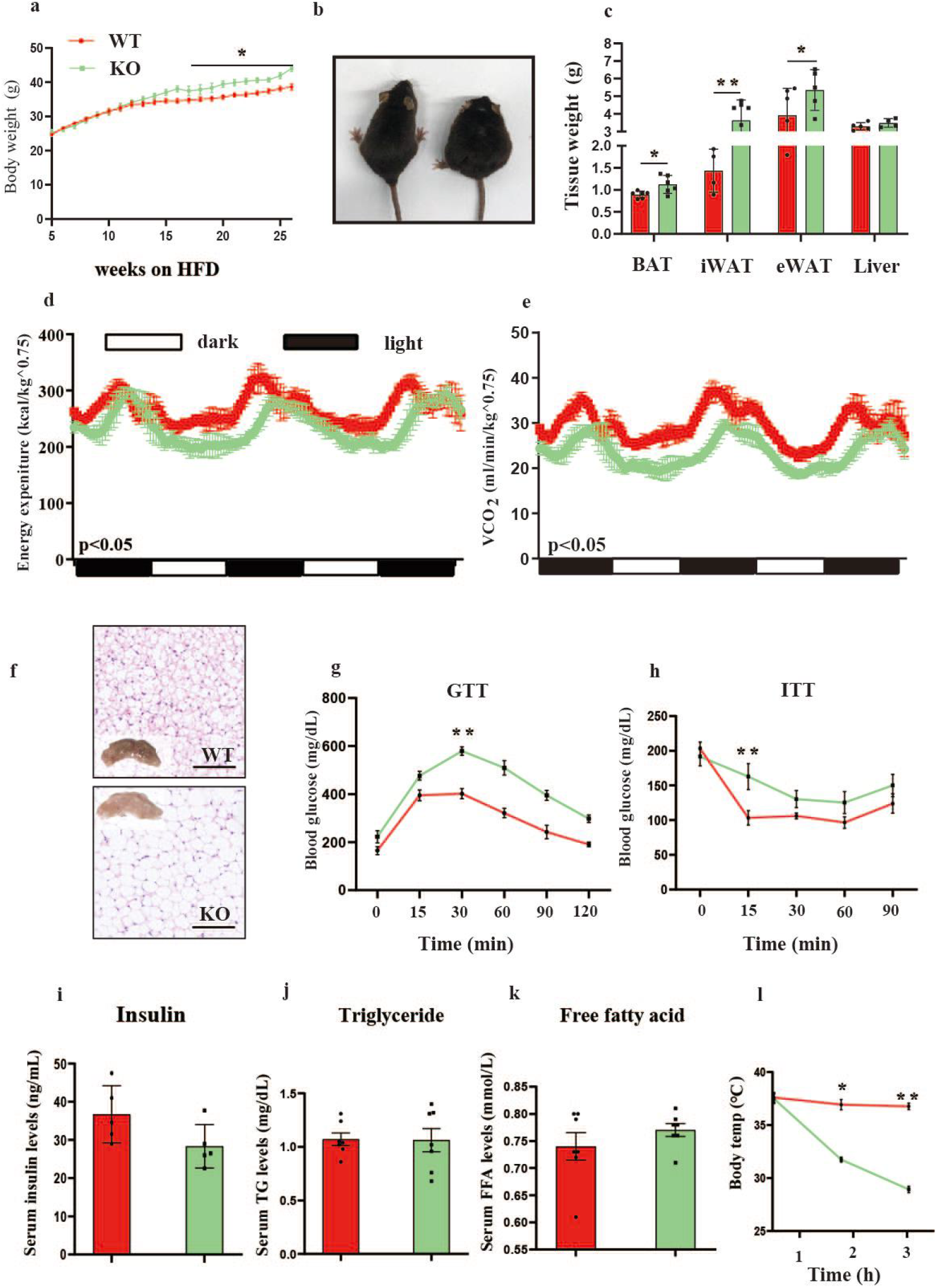
*Letmd1* deficiency leads to diet induced obesity and insulin resistance. (a) Whole body weight curves measured every week during 30 w with HFD-induced (n =7 for each group). (b) Representative image of WT and KO mice after 30 w HFD-induced. (c) tissue weight of BAT, iWAT, eWAT, Liver measured after 30 w HFD-induced (n = 7 for each group). (d-e) The energy expenditure and CO_2_ consumption rates in 30 w HFD-induced WT and KO mice at 22°C (n = 5 for each group) measured by CLAMS system. (f) Representative optical microscopy images of H&E staining BAT from HFD-fed WT and KO mice (Scale bar 200 μm). (g) The blood glucose levels with glucose injection (1 g/kg) measured by the glucometer as GTT for WT and KO mice fed with HFD (n = 6 for each group). (h) The blood glucose levels with insulin injection (1.5 U/kg) measured by the glucometer as ITT for WT and KO mice fed with HFD (n = 6 for each group). (i-k) Serum levels of Triacylglycerol (TG), Free Fatty Acids (FFA) and Insulin in WT and KO mice fed with HFD (n = 7 for each group). (l)The core temperature was tested by the thermo detector every 2 h of WT and KO mice fed with HFD in acute cold exposure (4°C) (n = 6-8 for each group). Data were mean ±SEM and each dot showed as one replicate. *P < 0.05, **P < 0.01, ***P < 0.001, Unpaired two-tailed Student’s t-tests was used in two sets of data.

In addition, after 4 h of cold exposure, the core body temperature of *Letmd1* KO mice dropped below 30°C (Fig. 5l).

### 3.6 Loss of Letmd1 leads to imbalanced mitochondrial fission and the calcium homeostasis

Mitochondrial remodeling plays an important role in BAT thermogenesis and metabolism^[39, 40]^. In the experiment of acute cold exposure, we found that the expression of dynamin-related GTPase (DRP1), a fission marker of mitochondria was decreased, meanwhile cold exposure increased the expression of key fusion protein MFN2 in BAT of *Letmd1* KO mice compared to the WT mice (Fig. 6a). Furthermore, we observed the elongated mitochondria in Letmd1 KO primary brown pre-adipocytes (Fig. 6b). The pre-adipocytes isolated from *Letmd1* KO mice had the reduced expression of TOMM20 in protein (Fig. 6c).

**Fig. 6.**
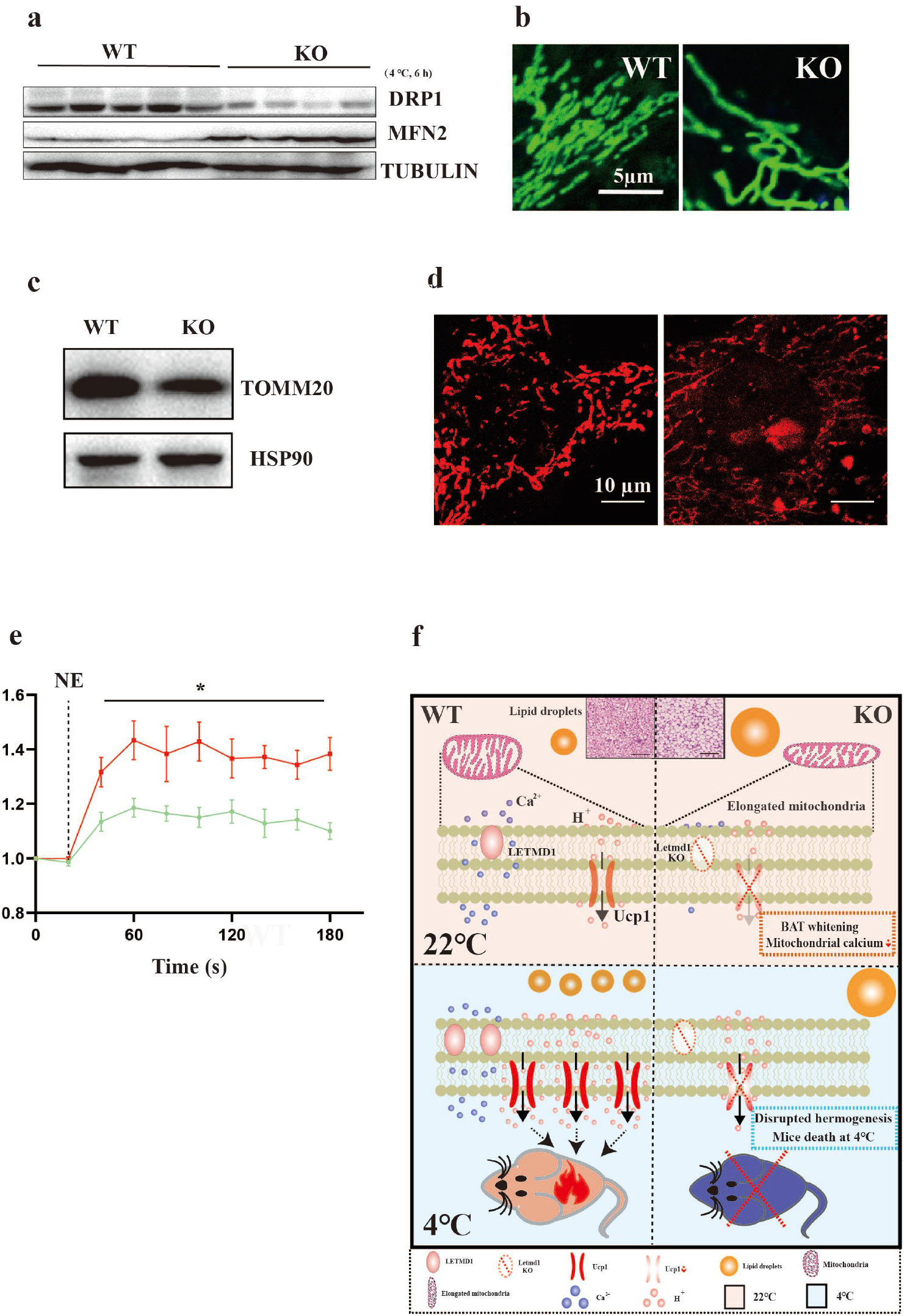
Loss of Letmd1 leads to imbalanced mitochondrial fission and the calcium homeostasis. (a) Immunoblotting measurements of DRP1 and MFN2 from BAT of WT and KO mice following 6 h at 4°C treatment (n = 4-5 in each group), α-TUBULIN was used as a loading control. (b) Representative microscopy images of mitochondrial morphology (green) with mitotracker staining in pre-adipocytes isolated from BAT of WT and KO mice. (c) Representative immunoblotting measurement of TOMM20 proteins in pre-adipocytes isolated from WT and KO mice, HSP90 was used as a loading control. (d) Representative microscopy images of mitochondrial Ca^2+^ (Red) with rhod-2 staining in brown adipocytes isolated from BAT of WT and KO mice. (e) NE-induced Ca^2+^ release in differentiated brown adipocytes isolated from WT and KO in a Ca^2+^-depleted medium. (f) Diagram models of the effects of Letmd1 KO mice, such as the BAT whitening, disruption of thermogenesis, cold-induced death, diet-induced obesity, hyperglycinemia and insulin resistance with the reduction of mitochondrial calcium ion and suppression fission of mitochondrion. Data were mean ±SEM and each dot showed as one replicate. *P < 0.05, **P < 0.01, ***P < 0.001, Unpaired two-tailed Student’s t-tests was used in two sets of data.

Recent studies have reported that calcium could regulate the morphology and the functions of mitochondria including the thermogenesis, fission and fusion^[41, 42]^. We found that the mitochondrial calcium concentration of brown adipocytes isolated from *Letmd1* KO was lower than that in WT (Fig. 6d and e). These results suggested that the loss of *Letmd1* might cause the imbalance of mitochondrial calcium homeostasis, which resulted in the abnormities of mitochondrial structures and functions, especially for the thermogenesis.

## 4 Discussion

*LETMD1* as an oncogene involved in the tumorigenesis of a variety of cancers has been extensively studied^[1, 6, 7]^. The high expression and the mitochondrial localization of LETMD1 in BAT indicated that LETMD1 might participant in BAT metabolism, whereas its precise function in BAT has not been addressed so far. Surprisingly, by generating and analyzing the phenotypes of the *Letmd1* KO mice, we found that the KO mice displayed multiple abnormalities related to metabolism, such as, BAT whitening, disruption of thermogenesis, cold-induced death, diet-induced obesity, hyperglycinemia and insulin resistance. Moreover, we demonstrated that the deletion of *Letmd1* in BAT destructed the mitochondrial calcium homeostasis, suppressed the mitochondrial fission, and disrupted UCP1-mediated BAT thermogenesis.

A growing number of studies suggest that LETMD1 is highly expressed in various malignant tumors tissues, such as cervical cancer, breast cancer, HCC, gastric cancer and colon cancer^[5–7, 9]^. Clinical studies have shown that the high expression of LETMD1 was related to the overexpression of breast cancer prognostic markers, such as human epidermal growth factor receptor 2 (HER2), and the presences of estrogen receptor (ER), progesterone receptor (PR) and p53 mutation^[6]^. Another study have demonstrated that the increased expression of LETMD1 could be a bio-marker for the assessment of postoperative prognostic for gastric cancer patients^[7]^. However, the functions of LETMD1 in metabolism are undefined. In this study, we uncovered that LETMD1 played an important role in metabolism and thermogenesis of BAT in mice.

The main function of BAT is thermoregulation with the uncoupling heat production to maintain body temperature of mammals, especially in cold environment^[28, 43]^. *UCP1* is the most important thermogenesis gene in BAT^[24, 44]^, and previous studies have showed that *Ucp1* deficiency destroyed thermogenesis, and the body temperature of 75% *Ucp1* KO mice dropped 10°C within 2 to 8 h^[44]^. Another study has demonstrated that *Aifm2* was required for cold-induced thermogenesis, and half of *Aifm2* KO mice died within 8 h at 4°C cold exposure^[27]^. Consistent with previous studies, we also found that the expression of UCP1 was drastically downregulated in *Letmd1* KO mice and the core body temperature dropped sharply. Unexpectedly, *Letmd1* KO mice died within 6 to 8 h at 4°C exposure, which implicated that deletion of *Letmd1* suspended the thermogenesis, and *Letmd1* was a crucial factor to regulate thermogenesis in BAT.

Mitochondrial dysfunctions suppress the metabolism and thermogenesis in BAT^[29]^. A study has shown that the peroxisomal biogenesis factor 16 *(Pex16)* deficiency inhibited the mitochondrial fission, decreased the mitochondrial copy number, and caused the mitochondrial dysfunction and in turn impaired the cold tolerance, decreased the energy expenditure, and increased the diet-induced obesity in mice^[45]^. Another study has reported that *Letm1* affected the expression level of UCP1 by regulating the homeostasis of mitochondrial calcium ions, the hydrogen ions, and the calcium ions mediated mitochondrial-nuclear crosstalk to regulate mitochondrial respiratory in BAT^[46]^. Recently, Lim’s study was the first one mentioned that knockdown of *LETMD1* decreased the ATP production and increased the intracellular ROS and calcium ions levels in macrophages^[10]^. Consistent with these findings, our data showed that deletion of *Letmd1* led to the decreases of mitochondrial copy numbers and the dysfunctions of mitochondria through destroying mitochondrial calcium homeostasis, However, different with Lim’s study, we found that Letmd1 KO decreased the concentration of mitochondrial calcium ions. As we showed in this study, we did not observe the phenotypical changes in liver, iWAT and eWAT caused by *Letmd1* deletion, which might depend on the higher expression and intense metabolic capacity of mitochondria in BAT.

Energy expenditure reductions, obesity and hyperglycemia are the signs of dysfunctions of BAT associated with mitochondrial dysfunctions^[47]^. A study has shown that the deletion of transient receptor potential cation channel subfamily V member 2 *(Trpv2)* impaired BAT thermogenesis and increased the body weight and fat with HFD-diet treatment^[48]^. Consistent with these studies, we found that *Letmd1* KO mice showed the excessive obesity, glucose intolerance and insulin resistance under the HFD condition. The capacity to tolerate the cold of the HFD-induced *Letmd1* KO mice was deteriorated more severe than that of animal with the Chow diet. This indicated that the mitochondrial heat producing capacity of the obese *Letmd1* KO mice was further damaged.

The clinical evidences of LETMD1 in obesity and metabolism remain unclear, and whether LETMD1 is a mitochondrial calcium transportation protein is still unknown. In future study, we will future analyze the clinical data of metabolic diseases for exploring the relationship between LETMD1 and metabolic disorders. We will also explore how LETMD1 regulates the ions transportation and BAT metabolism with the LETMD1 overexpression and mutation animal models.

In summary, we uncovered a novel molecular mechanism of LETMD1 in metabolism and BAT thermogenesis (Fig. 6f). Deletion of *Letmd1* caused the defectives of mitochondrial calcium ion homeostasis in BAT, leading to abnormal mitochondrial structures and functions, and severely impairing heat generation and BAT whitening, which in turn resulting the disruption of thermogenesis, cold induced death, diet-induced obesity, hyperglycinemia and insulin resistance. Our data suggest that Letmd1 may be a therapeutic target for clinical treatment of metabolic diseases and obesity.

## Conflict of interest

The authors declare that they have no conflict of interest.

## Acknowledgments

We thank for the instrument and equipment support provided by the platform of Institute of Nutrition and Health, China Agricultural University.

## Funding

This work was supported by the National Key Research and Development Project (2018YFC1004702); Natural Science Foundation of Beijing (20G10613); Natural Science Foundation of China (NSFC31970802); The Project for Extramural Scientists of State Key Laboratory of Agrobiotechnology (2018SKLAB6-19).

## Author contributions

Xiangdong Li signed the project, guided experiments, and analyzed data. Runjie Song, Yaqi Du, Peng Li and Xiangdong Li interpreted the data. Runjie song, Yaqi Du and Peng Li conducted experiments and collected data. Huijiao Liu, Lijun Zhou and Han Zheng assisted animal studies. Huijiao Liu and Xiaohui Lu analyzed data. Shenghong Wang made revisions to the manuscript. Fazheng Ren and Xiru Li provided guidance for experiments. All authors approved the final content.

## Notes

### Competing Interest Statement

The authors have declared no competing interest.

