## Supplemental fig for "Deletion of *Letmd1* leads to the disruption of mitochondrial function in brown adipose tissue"

**Fig. S1** (a-d) The $O_2$ consumption rates, Respiratory Quotient and food intake in 8-w-old WT and KO mice fed chow diet at 22°C measured by CLAMS system (n = 5 for each group). (e-f) Representative optical microscopy images of H&E staining iWAT, eWAT and Liver from WT and KO mice fed chow diet at 22°C (Scale bar 200 μm). (g) Immunoblotting measurement of TOMM20 in different mice tissues (n = 3), α-TUBULIN was used as a loading control. (h-i) The $O_2$ consumption rates, respiratory quotient and food intake in 30 w HFD-induced WT and KO mice at 22°C measured by CLAMS system (n = 5 for each group). Data were mean ± SEM and each dot showed as one replicate. *$P < 0.05$, **$P < 0.01$, ***$P < 0.001$, Unpaired two-tailed Student's t-tests was used in two sets of data.

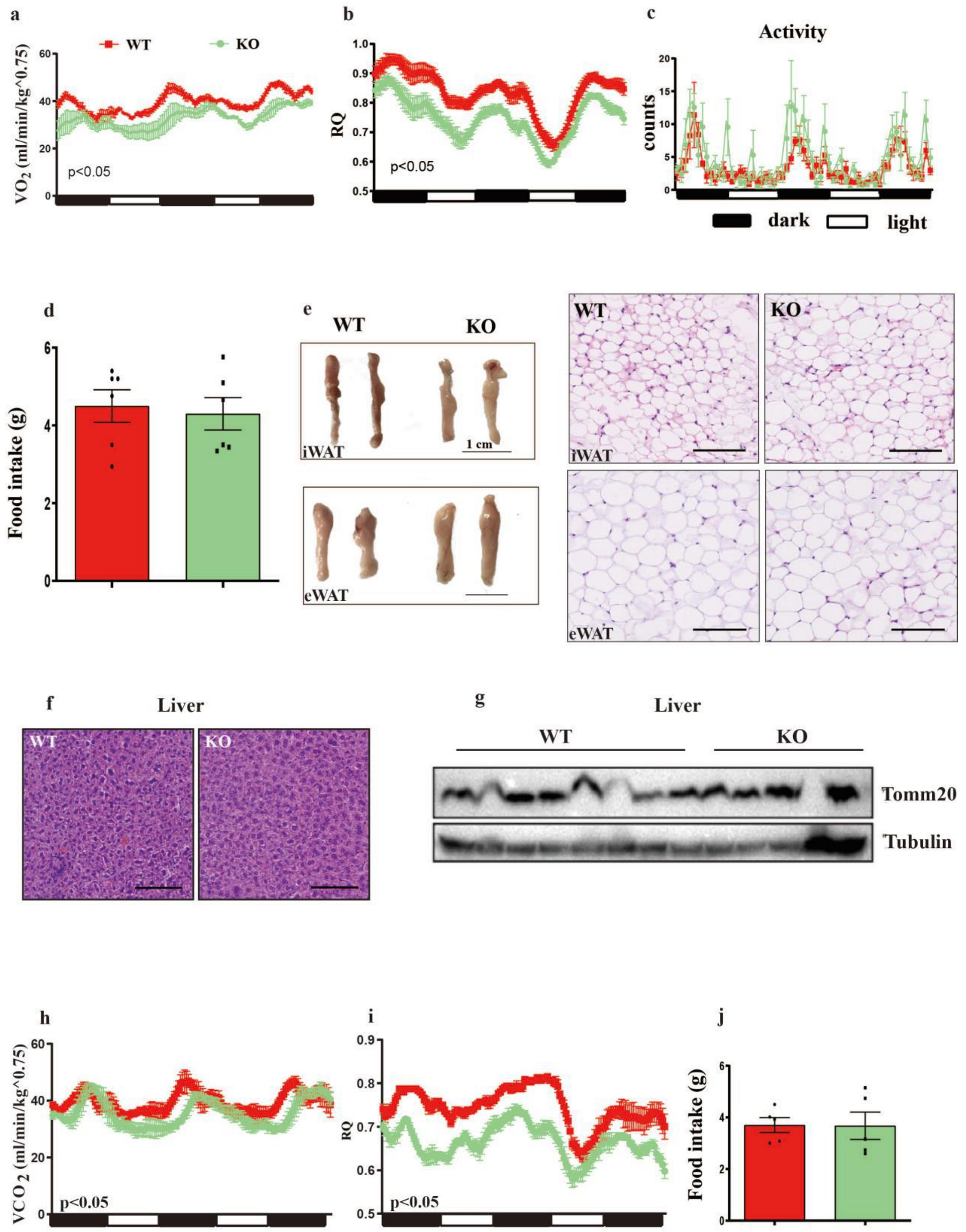

a
WT
KO
VO2 (ml/min/kg^0.75)
p<0.05
b
RQ
p<0.05
c
Activity
counts
dark
light
d
Food intake (g)
e
WT
KO
iWAT
eWAT
1 cm
f
Liver
WT
KO
g
Liver
WT
KO
Tomm20
Tubulin
h
VCO 2 (ml/min/kg^0.75)
p<0.05
i
RQ
p<0.05
j
Food intake (g)
