## Supplementary table for "Deletion of *Letmd1* leads to the disruption of mitochondrial function in brown adipose tissue"

**Supplementary Table 1. Primers list for RT-qPCR**

| **Gene** | **Forward primer (5' - 3')** | **Reverse primer (5' - 3')** |
|---|---|---|
| *Ucp1* | CACTCAGGATTGGCCTCTACG | GGGGTTTGATCCCATGCAGA |
| *mitoDNA* | TTAAGACACCTTGCCTAGCCACAC | CGGTGGCTGGCACGAAATT |
| *Dio2* | GATGCTCCCAATTCCAGTGT | TGAACCAAAGTTGACCACCA |
| *β-ACTIN* | GGCTGTATTCCCCTCCATCG | CCAGTTGGTAACAATGCCATGT |
| *Fasn* | AGGATATGGAGAGGGCTGGT | ACCCAAGCATCATTTTCGTC |
| *Pgc1α* | CCCTGCCATTGTTAAGACC | TGCTGCTGTTCCTGTTTTC |
| *Cidea* | TGACATTCATGGGATTGCAGAC | GGCCAGTTGTGATGACTAAGAC |
| *Elovl3* | TTCTCACGCGGGTTAAAAATGG | GAGCAACAGATAGACGACCAC |
| *Atp5a* | TTTGCCCAGTTTGGTTCTGAT | CCCGTACACCCGCATAGATAA |
| *Uqcrc2* | TGGCTCTGGTTGGACTTGGT | TTTCACCTCCACGGTATTTGG |
| *Sdhb* | CTGTCGAGGGGCACAGAC | CAACACCATAGGTCCGCACT |
| *Ndufb8* | GGCACGTGTTCCCTTCCTAC | CCGCTCCAGGTACAGATTATTGT |
| *Cox8b* | GAACCATGAAGCCAACGACT | GCGAAGTTCACAGTGGTTCC |
| *HSL* | GAGACACCAGCCAACGGATAC | TTTTGCGGTTAGAAGCCACAT |
| *Cpt2* | GGCCACCAACTTGACTGTTT | GAAGGAACAAAGCGGATGAG |
| *Glut1* | TCAACACGGCCTTCACTG | CACGATGCTCAGATAGGACATC |
| *HK2* | TGATCGCCTGCTTATTCACGG | AACCGCCTAGAAATCTCCAGA |
| *Mtco1* | ACACATGAGCAAAAGCCCAC | AGTCTGAGTAGCGTCGTGGT |
| *PPARγ* | CCGTAGAAGCCGTGCAAGAG | GGAGGCCAGCATCGTGTAGA |
| *NucDNA* | ATGACGATATCGCTGCGCTG | TCACTTACCTGGTGCCTAGGGC |
| *Letmd1* | CGGGAGATGGAGCATTTGAGACAG | GGCAAAGGGTGGAATGGAGATGAG |
| *Pgc1α* | CCC TGC CAT TGT TAA GAC C | TGC TGC TGT TCC TGT TTT C |

**Supplementary Table 2. Antibody list**

| **Antibody** | **Company** |
|---|---|
| UCP1 | Abcam, USA |
| LETMD1 | Abcam, USA |
| SCA-1-APC | Miltenyi Biotec, Germany |
| CD11B-FITC | Miltenyi Biotec, Germany |
| CD45-PE | Miltenyi Biotec, Germany |
| HSP90 | Proteintech, USA |
| TOMM20 | Proteintech, USA |
| A-TUBULIN | Proteintech, USA |
| MFN2 | Proteintech, USA |
| DRP1 | Proteintech, USA |
